# Unique gut microbiome signatures among adult patients with moderate to severe atopic dermatitis in southern Chinese

**DOI:** 10.1101/2022.05.14.491964

**Authors:** Yiwei Wang, Jinpao Hou, Joseph Chi-Ching Tsui, Lin Wang, Junwei Zhou, Un Kei Chan, Claudia Jun Yi Lo, Pui Ling Kella Siu, Steven King Fan Loo, Stephen Kwok Wing Tsui

**Author notes:** **Corresponding author:** Stephen Kwok Wing TSUI, Steven King Fan LOO.

## Abstract

Imbalance of the immune system caused by alterations of gut microbiome is considered to be a critical factor in the pathogenesis of infant eczema but the exact role of the gut microbiome in adult atopic dermatitis (AD) patients remains to be clarified. To investigate the differences of the gut microbiome between adult AD patients and healthy individuals, stool samples of 234 adults, containing 104 AD patients and 130 healthy subjects were collected for amplicon sequencing. Altered structure and metabolic dysfunctions of the gut microbiome were identified in adult AD patients. Our results illustrated that the adult AD patients were more likely to have allergies, particularly non-food allergies. And the gut microbiome composition of the AD and normal groups were considerably different. Besides, *Romboutsia and Clostridium_sensu_stricto_1* was enriched in the normal group, whereas *Blautia, Butyricicoccus, Lachnoclostridium, Eubacterium_hallii_group, Erysipelatoclostridium, Megasphaera, Oscillibacter, Flavonifractor* were dominated in the AD group. Moreover, purine nucleotide degradation pathways were significantly enriched in the AD group and the enrichment of proteinogenic amino acid biosynthesis pathways was found in the normal group. This study provides insights into new therapeutic strategies targeting the gut microbiome for AD and evidence for the involvement of gut-skin axis in AD patients.

## INTRODUCTION

Atopic dermatitis (AD), also commonly referred to as eczema, is one of the most common skin diseases, characterized by recurrent chronic eczematous lesions, occurring with dry skin, pruritus, and obvious itching(Langan et al., 2020). In the past decade, the global incidence of atopic dermatitis has been increasing, and the affected population involves all age groups ranging from infants to the elderly. This seriously influences the patients’ quality of life and brings a huge burden to healthcare resources(Langan et al., 2020). AD patients usually may also be accompanied by other atopic diseases, such as allergic asthma and allergic rhinoconjunctivitis(Caubet and Eigenmann, 2010). The pathogenesis of atopic dermatitis is not yet clear. It is generally believed that AD may be related to genetics, environmental factors, immune abnormalities, and abnormal skin barrier function. It is the interaction between genetic factors and environmental factors that mediate the occurrence and development of AD through immune pathways(Lee et al., 2018).

The gut microbiome is renowned as the second human genome, the homeostasis of which has been evidenced to have a beneficial effect on maintaining human health and is commonly affected by many factors, such as age, gender, diet, mood, and health status(Jackson et al., 2018, Zhu et al., 2010). The gut flora of adults is proved to be more complex than that of infants(Galkin et al., 2020). Imbalance of the gut microbiome in early childhood precedes the onset of atopic dermatitis(Lee et al., 2018). Numerous studies have focused on the characterization of the gut microbiome in AD infants or children because of the strong impact of the gut microbiome on the development and maintenance of the immune system in early life(Abrahamsson et al., 2012, Hong Pei-Ying et al., 2010, Ismail et al., 2012, Laursen Martin Frederik et al., 2015, Lee Eun et al., 2016, West et al., 2015).In the intestines of AD infants, the relative abundance of *Bifidobacterium, Enterococcus, Clostridium, Lactobacillus paracasei*, and *Ruminococcaceae* decrease(Gore et al., 2008, Hong Pei-Ying et al., 2010, West et al., 2015). In contrast, gut colonization with *Staphylococcus, Clostridia*, and *Feacalibacterium prausnitzii* is more prevalent in AD infants(Penders et al., 2013, Song et al., 2016). However, few studies have clarified the role of the gut microbiome in adult AD patients.

In this study, we performed amplicon sequencing and revealed the differences in gut microbiome composition, biodiversity, and functional profiling using amplicon sequencing between healthy adults and AD patients from a large Hong Kong cohort.

## RESULTS

### Characteristics of participants

A total of 104 AD patients and 130 healthy subjects were recruited in this study. Baseline characteristics of participants were summarized in Table 1. A higher proportion of participants with a medical history of allergy (p = 0.0002), non-food allergy (p = 0.0301) especially, were observed among the AD group.

**Table 1.** Characteristics in AD group and normal group

| Parameters |  | AD (n = 104) |  | Normal (n = 130) |  | *p* value<sup>1</sup> |
| --- | --- | --- | --- | --- | --- | --- |
|  |  | *mean ± sd* | n | *mean ± sd* | n |  |
| Age |  | 44.3 ± 15.1 |  | 47.7 ± 15.1 |  | 0.2155 |
| Sex | Female |  | 63 |  | 78 | 1.0000 |
|  | Male |  | 41 |  | 52 |  |
| Allergy<sup>2</sup> | Yes |  | 31 |  | 10 | **0.0002** |
|  | No |  | 32 |  | 50 |  |
|  | Not sure |  | 16 |  | 8 |  |
| Food allergy<sup>2</sup> | Yes |  | 12 |  | 5 | 0.1279 |
|  | No |  | 65 |  | 63 |  |
|  | Not sure |  | 2 |  | 0 |  |
| Non-food allergy<sup>2</sup> | Yes |  | 17 |  | 5 | **0.0301** |
|  | No |  | 62 |  | 63 |  |
| Diarrhea<sup>2</sup> | Yes |  | 17 |  | 18 | 0.6110 |
|  | No |  | 62 |  | 50 |  |
| Constipation<sup>2</sup> | Yes |  | 26 |  | 30 | 0.2207 |
|  | No |  | 53 |  | 38 |  |
| BMI<sup>2</sup> |  | 23.3 ± 3.7 |  | 23.1 ± 3.8 |  | 0.7090 |
| Overweight (BMI > 25)<sup>2</sup> | Yes |  | 21 |  | 19 | 0.9828 |
|  | No |  | 52 |  | 44 |  |
|  | Not sure |  | 6 |  | 5 |  |

<sup>1</sup>*p* value was calculated using Chi-square test and Wilcoxon rank-sum test
<sup>2</sup> 147 of 243 participants provided the clinical information of allergy history, prevalence of diarrhea and constipation, BMI (both height and weight) for subsequent patient characteristics analysis.

### Distinct gut microbiome composition between AD patients and healthy subjects

To dissect potential dysbiosis of gut bacterial communities among AD patients, we compared the alpha and beta diversity between AD group and normal groups. Although no significant difference (Shannon diversity index, *p* = 0.285) in alpha diversity, including richness and evenness of the gut microbiome was obtained between the AD groups and the normal group (Figure 1AB, Table S1). Beta diversity analysis results demonstrated that gut microbiome of AD group was immensely distinct from that of normal group based on Bray-Curtis (*p* = 0.001), Jaccard (*p* = 0.001) and unweighted-UniFrac (*p* = 0.025) distance metrics (Figure 1C, Table S2). Furthermore, statistically significant difference in beta diversity based on Bray-Curtis (*p* = 0.001), Jaccard (*p* = 0.001), and unweighted-UniFrac (*p* = 0.031) were obtained between normal and severe AD groups, as well as normal and mild AD groups (Figure 1D-F, Table S2). Collectively, these findings indicate differences in the gut microbial community structure between patients with AD and healthy individuals.

**Figure 1.**
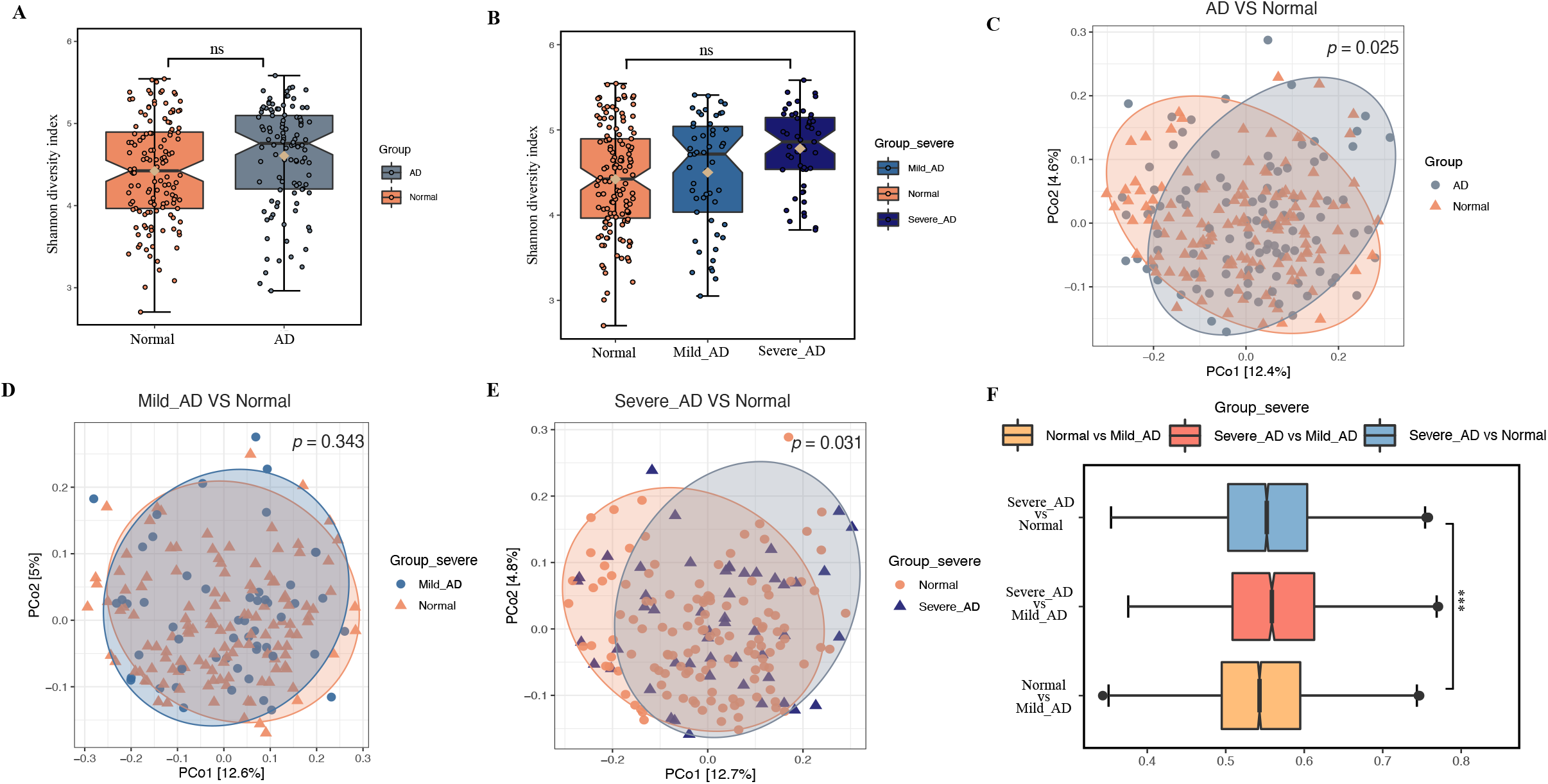
Alpha diversity and beta diversity analyses of the gut microbiome between patients with AD and healthy subjects. (A), (B) illustrated the comparison of Shannon diversity index across groups. Abbreviations: ns, no significant difference was calculated by the Kruskal-Wallis test. (C-F) Beta diversity analysis across groups. Principal Coordinate Analysis (PCoA) plot based on unweighted UniFrac distances, coloured by groups - (C) AD VS Normal, (D) Mild_AD VS Normal, (E) Severe_AD VS Normal. *P* value was calculated by PERMANOVA test with permutation = 999. (F) Comparison of unweighted UniFrac distances across normal, Mild_AD, and Severe_AD groups showing the dissimilarity of the gut bacterial communities. *P* value was calculated using the Wilcoxon rank-sum test. *** *p*<0.001.

### Gut microbial composition in healthy individuals and AD patients

At the phylum level, a total of 15 phyla, including 12 from the kingdom of bacteria and 3 from archaea, were detected which constituted the gut microbiome of both patients with AD and healthy individuals (Figure2A, Table S3), and the top six most abundant phyla accounted for over 99% of sequences in the dataset (Figure 2B). In both AD and normal group, *Firmicutes* (AD, 57.4%; normal 58.2%) was dominant, followed by *Bacteroidota* (AD, 32.4%; normal 32.1%), *Actinobacteriota* (AD, 6.2%; normal, 5.5%), *Proteobacteria* (AD, 2.7%; normal, 2.4%), *Desulfobacterota* (AD, 0.5%; normal, 0.7%) and *Fusobacteriota* (AD, 0.3%; normal, 0.5%; Figure 2B, Table S3). Besides, by conducting ANCOM analysis, the gut microbial community of the AD group demonstrated a similar composition with the normal group at the phylum level (Table S3). Additionally, no significant difference (AD VS normal, *p* = 0.7566; Mild AD VS Severe AD VS normal, *p* = 0.5748) of *Firmicutes* to *Bacteroidetes* (F/B) ratio was detected across groups (Figure S1). At the genus level, the topmost abundant were *Bacteroides, Prevotella, Blautia, Faecalibacterium*, and *Bifidobacterium* (Figure 2C) in both groups. Furthermore, only the relative abundance of *Clostridium_sensu_stricto_1* genus in the normal group was significantly higher than that in the AD group (W=218), using ANCOM (Figure2D).

**Figure 2.**
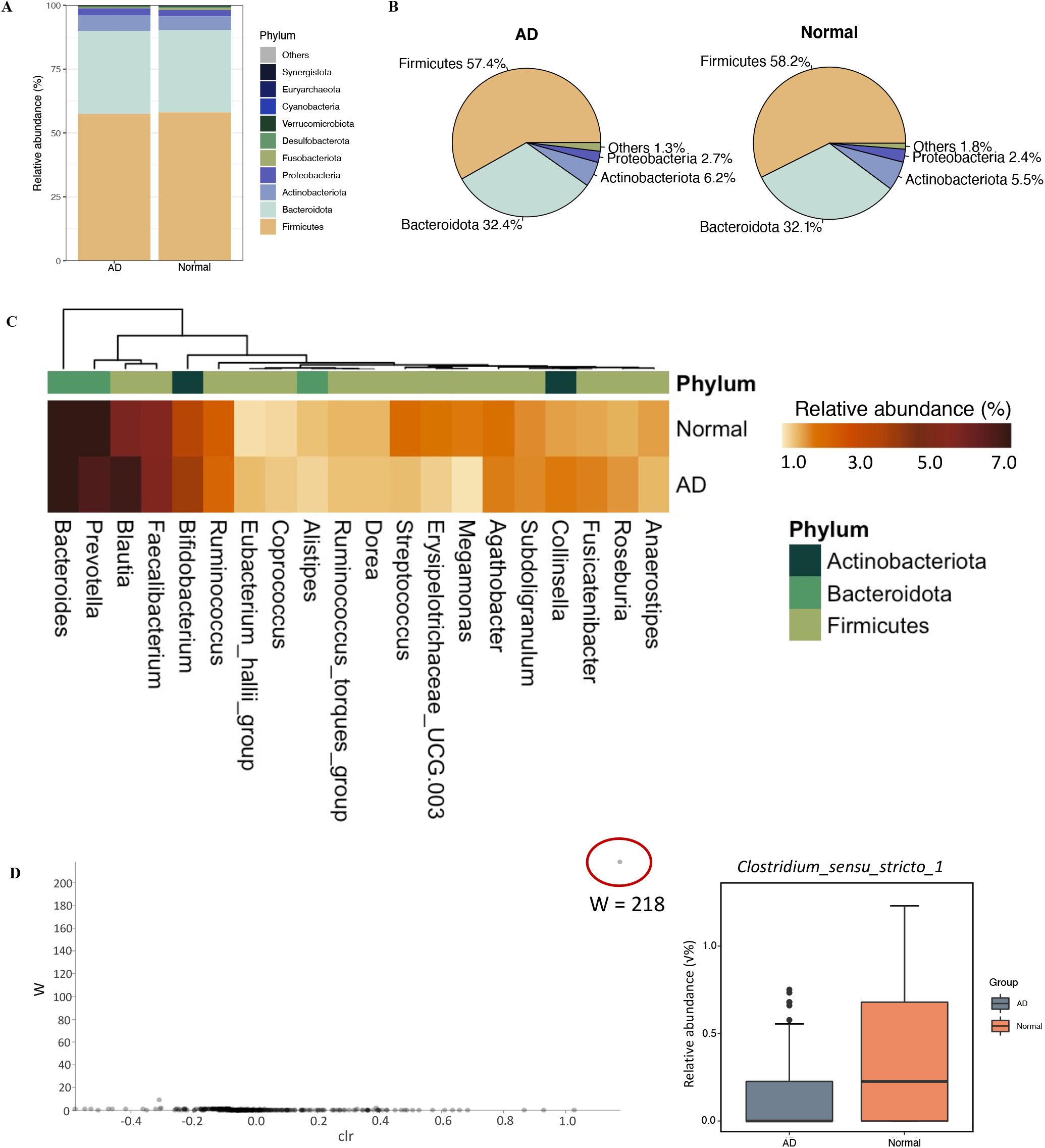
Taxonomic profiling of gut bacterial composition and differential abundant test results using ANCOM between AD and normal groups. (A) Bar plot and (B) pie charts of bacterial composition at the phylum level across groups. (C) Top 20 genera in the gut microbiome of AD and normal groups. (D) Differential abundant genus using ANCOM and boxplot of the relative abundance of Clostridium_sensu_stricto_1. Abbreviations: clr, centered log-ratio transformation on the relative abundance of each SV.

### Identification of gut microbial signatures to differentiate between AD and healthy subjects

A total of 13 genera were identified as gut bacterial signatures at the genus level between AD patients and healthy individuals using the method of LEfSe (Figure 3, Table S4). Nine genera, including *Blautia, Butyricicoccus, Lachncoclostridium, Eubacterium_hallii_group, Erysipelatoclostridium, Megasphaera, Oscillibacter, Flavonifractor*, and unclassified genera from the bacteria family of *Oscillospiraceae* were significantly enriched in the AD group. The other 4 genera, containing *Romboutsia, Clostridium_sensu_stricto_1* and unclassified genera from two bacteria families *Butyricicoccaceae* and *Erysipelotrichaceae* were considerably enriched in the normal group (Figure 3A, Table S4). Additionally, the relative abundance of *Blautia* and *Butyricoccus* was significantly positively correlated with the severity of AD. While as the severity of AD deepens, the relative abundance of two genera, *Romboutsia* as well as *Clostridium_sensu_strico_1*, and one family *Erysipelotrichaceae* decreases significantly (Figure3B). Additionally, the enrichment of *Bacteroides, Butyricicoccus*, and *Lachnospiraceae_UCG_008* was notably obtained in the Mild_AD group, while the relative abundance of *Blautia* in the Severe_AD group was significantly higher than that in normal and Mild_AD groups (Figure S2).

**Figure 3.**
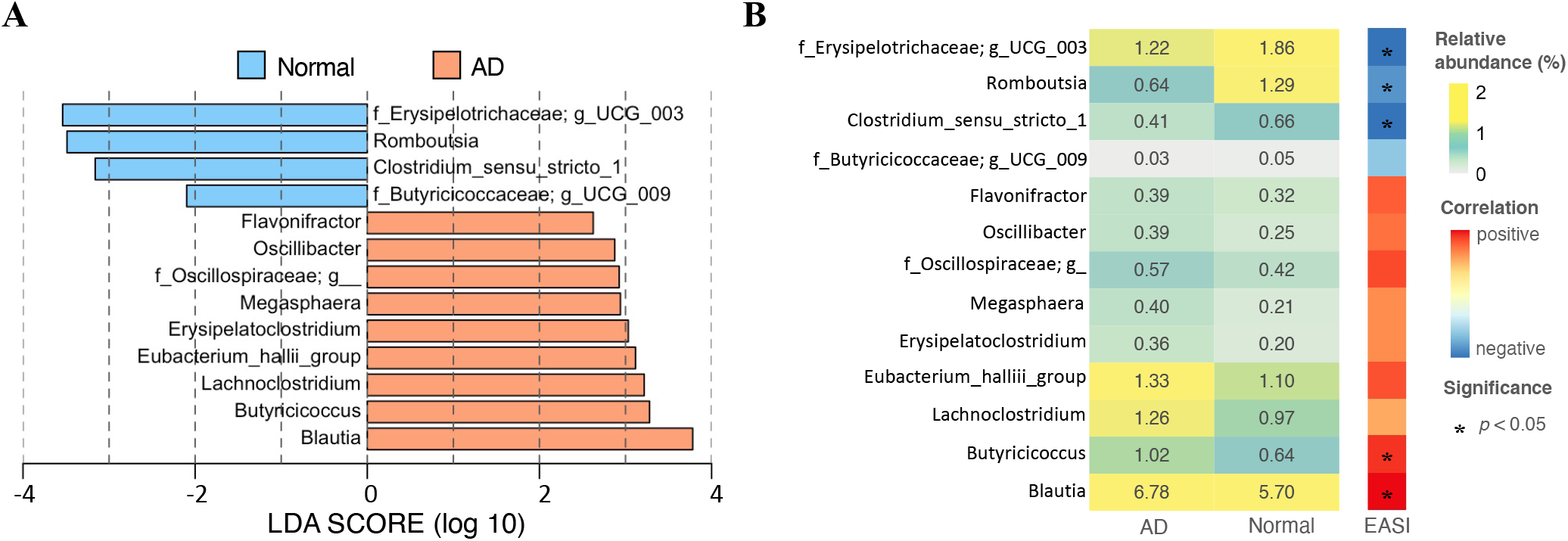
Gut microbial signatures selected by LEfSe analysis and relative abundance of gut microbial biomarkers across groups. (A) Bar plot of LEfSe analysis result (B) Heatmap displayed the relative abundance of signatures identified using LEfSe analysis in AD and normal groups and correlation between the EASI score and the relative abundance of signatures.

### Significantly altered MetaCyc pathways of the gut microbiome associated with AD

We next performed functional prediction analysis on the gut metagenome using PICRUSt2 and LEfSe algorithms. This identified the difference in the function of the gut microbiome between AD patients and healthy subjects. Distinct MetaCyc pathways differences between AD and normal groups were calculated by LEfSe analysis on PICRUSt2 output (Figure 4, Table S5). Specifically, dTDP-N-acetylthomosamine biosynthesis (PWY-7315), D-fructuronate degradation (PWY-7242), purine nucleobases degradation I (anaerobic) (P164-PWY), formaldehyde assimilation II (RuMP Cycle) (PWY-1861), superpathway of glycolysis and Entner-Doudoroff (GLYCOLYSIS-E-D), superpathway of N-acetylneuraminate degradation (P441-PWY), methanogenesis from acetate (METH-ACETATE-PWY), guanosine nucleotides degradation III (PWY-6608), purine nucleotides degradation II (aerobic) (PWY-6353), adenosine nucleotides degradation II (SALVADEHYPOX-PWY) and palmitate biosynthesis II (bacteria and plants) (PWY-5971) were enriched in AD patients. In contrast, urea cycle (PWY-4984), nitrate reduction VI (assimilatory) (PWY490-3), GDP-mannose biosynthesis (PWY-5659), glycolysis I (from glucose 6-phosphate) (GLYCOLYSIS), L-lysine biosynthesis II (PWY-2941), superpathway of L-tyrosine biosynthesis (PWY-6630), as well as superpathway of L-phenylalanine biosynthesis (PWY-6628) were identified to be enriched in the normal group (Figure 4A, Table S5). Interestingly, most functional pathways (8/11) enriched in the AD group were assigned to degradation/utilization/assimilation superclass, of which 4 pathways were affiliated to purine nucleotide degradation class. Among the pathways enriched in the normal group, more than half (4/7) of the pathways were subjected to the superclass of biosynthesis, and 3 pathways belong to the class of proteinogenic amino acid biosynthesis class (Figure 4A). In addition, we also compared the differences in the functional potential of the gut microbiome between Mild_AD and Severe_AD patients. Purine nucleotides degradation II (aerobic) (PWY-6353) and adenosine nucleotides degradation II (SALVADEHYPOX-PWY), belonging to purine nucleotides degradation were found to be enriched in the Severe_AD group. By contrast, pyridoxal 5’-phosphate biosynthesis I (PRYIDOXSYN-PWY) and superpathway of pyridoxal 5’-phosphate biosynthesis and salvage (PWY0-845) pertaining to vitamin B6 biosynthesis were most abundant in the Mild_AD group (Figure 4B, Table S5).

**Figure 4.**
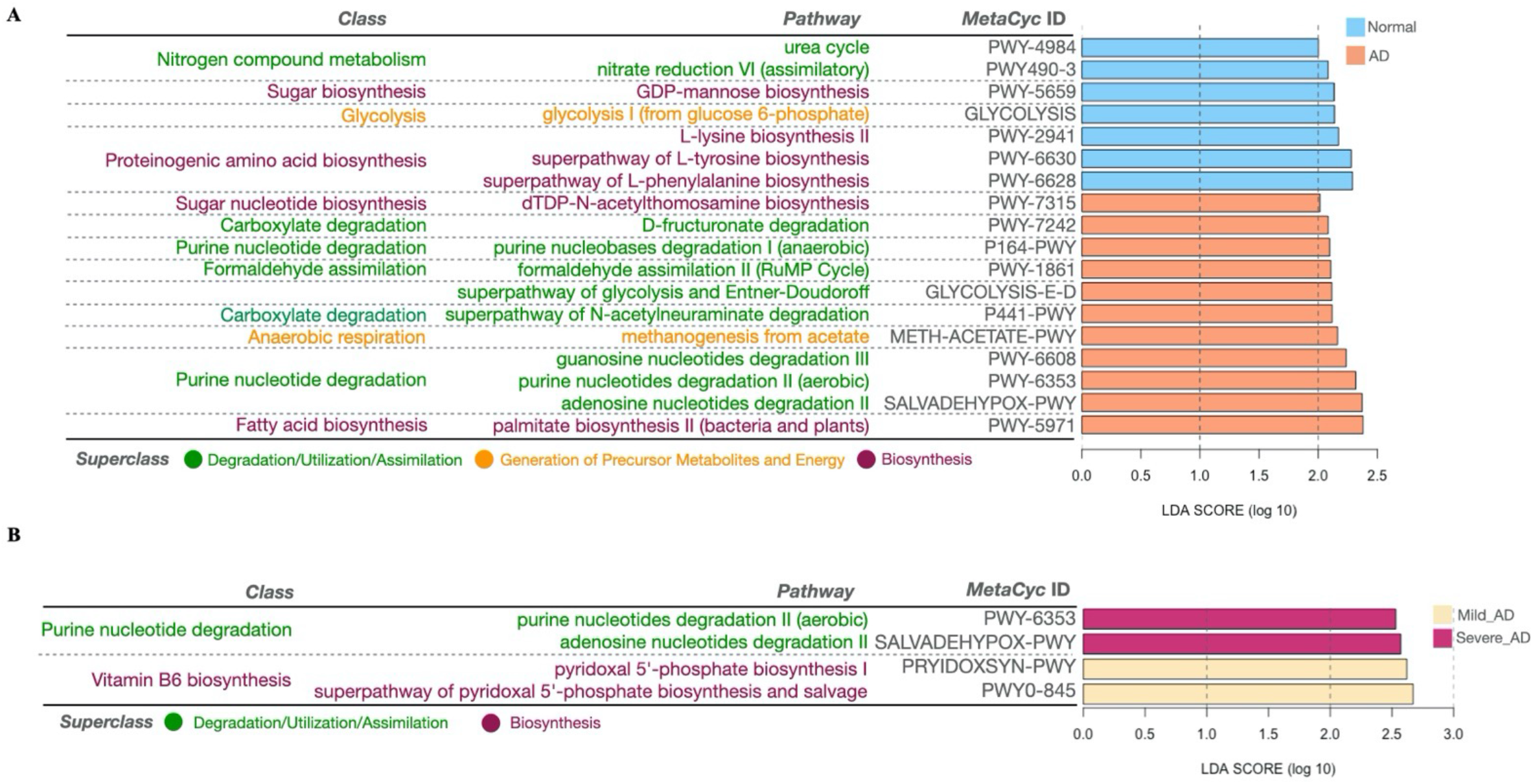
Differentially enriched predicted functional pathways across groups using PICRUSt2 and LEfSe analysis. The bar chart in (A) showed the LDA score (log10) of the enriched pathways between the normal and AD groups. The bar chart in (B) displayed the LDA scores (log10) of the enriched pathways in the Mild_AD and Severe_AD groups. Two levels of annotations of each pathway were labeled, and the color of the pathway indicated the superclass to which it belongs. Green represented the superclass of degradation/utilization/assimilation; Purple represented the superclass of biosynthesis; Orange represented the superclass of generation of precursor metabolites and energy.

## DISCUSSION

To the best of our knowledge, this study is currently the first large-scale community-based gut microbiome study on adult atopic dermatitis. We evaluated the association between the incidence of adult AD and age, gender, BMI, the prevalence of allergies, gastrointestinal symptoms. We also performed amplicon sequencing to investigate the characteristics of adult AD patients’ gut microbiome from several aspects including gut microbial composition, biodiversity, and functional potential.

Our results support previous studies indicating that the incidence of allergy among AD patients is higher than that of healthy individuals. One previous study reported that nearly 80% of AD children will develop asthma or allergic rhinitis(Eichenfield et al., 2003). Several lines of evidence suggested that AD patients are more sensitive to aeroallergens and food allergens, which play an important pathogenetic role in the development of AD(Caubet and Eigenmann, 2010, Eller et al., 2009, Schäfer, 2008). In our study, AD patients have higher rates of non-food allergies compared with the healthy subjects, which is consistent with the previous conclusion that adults and children over 5 years old appeared to be more sensitive to non-food allergens such as house dust mite(Fuiano et al., 2010, Pónyai et al., 2008). However, we failed to obtain a difference in the prevalence of food allergy between AD and normal groups. Actually, associations between AD and food allergy remain controversial. Eller E et al. and Hon KL et al. reported that infants and toddlers with AD are more prone to food allergies(Eller et al., 2009, Hon et al., 2008). Moreover, among AD patients, the rate of food allergy occurrence is ranging from 30% to 80%, depending on the population, however, the actual incidence of confirmed food allergies is much lower(Eller et al., 2009, Hill et al., 2004, Kvenshagen et al., 2009). Although Carol C et al. reported that children with atopic eczema had a higher incidence of diarrhea, especially in those with diffuse eczema, we did not observe differences in the frequency of gastrointestinal symptoms between adult AD patients and healthy individuals. Consistent with our results, their study also failed to obtain an increase in constipation in AD children(Caffarelli et al., 1998). These inconsistent results may be attributable to differences in age, geographical locations, and different scales of assessing gastrointestinal symptoms.

Previous research conclusions on alpha diversity of the gut microbiome in AD patients were controversial. In our study, we found that the gut microbial community of adult AD patients displayed a similar alpha diversity. Similarly, 5 studies identified no significant differences in the alpha diversity of gut microbiome in AD infants or children compared with healthy control counterparts(Bisgaard et al., 2011, Gore et al., 2008, Hong P. Y. et al., 2010, Laursen M. F. et al., 2015, Lee E. et al., 2016). Lee and colleagues reported a higher alpha diversity in infant atopic eczema(Lee Eun et al., 2016). In contrast, 3 other studies indicated that the alpha diversity of infants and young toddlers with eczema or at high risk for AD decreased compared to controls(Abrahamsson et al., 2012, Ismail et al., 2012, Wang et al., 2008).

We confirmed the difference in the composition of the intestinal bacteria of AD and normal groups and pointed out the potential microbial signatures. Our results demonstrated that compared to the normal group, the relative abundance of two bacterial genera, *Clostridium_sensu_stricto_1* and *Romboutsia* were reduced in AD patients. *Clostridium_sensu_stricto_1* was the only species with differential abundance test that was consistently obtained by the two algorithms of ANCOM and LEfSe. *Clostridium_sensu_stricto_1*, containing 160 species(Gupta and Gao, 2009) includes *Clostridium butyricum*, which has a strong ability to produce butyric acid, a type of short-chain fatty acids (SCFAs)(Cassir et al., 2016). Similarly, *Romboutsia sedimentorum* in the genus *Romboutsia* is evidenced to produce short-chain fatty acids such as acetic acid, isobutyric acid, and isovaleric acid by fermenting glucose(Wang et al., 2015). Short-chain fatty acids, which play an important role in the gut-skin axis, can stamp out the immune responses by inhibiting the cytokine production, migration, proliferation, and adhesion of inflammatory cells. SCFAs can also regulate the apoptosis and activation of immune cells by deactivation of NF-κB signaling pathways and inhibiting histone deacetylase, which promotes the cell proliferation that regulates various skin physiological functions, for example, regulating the differentiation of hair follicle stem cell and wound healing(Salem et al., 2018). Moreover, SCFAs can provide energy for the intestinal epithelium, and improve the integrity of the intestinal epithelial barrier to prevent microorganisms and toxins from entering the body fluid circulation to trigger Th2 immunity, and further resulting in disturbance of the skin homostatis(O’Neill et al., 2016). Additionally, the relative abundance of two bacterial families, including *Erysipelotrichaceae* as well as *Butyricicoccaceae*, were evidenced to enrich in normal group. Past studies have shown that the relative abundance of *Butyricicoccaceae* was negatively correlated with diabetes(Huang et al., 2021). Our results also suggested that the relative abundance of *Blautia* and *Butyricicoccus* in AD patients was greater than those in the normal group. Fang et al. observed that *Blautia* was significantly enriched before the administration of probiotics in AD patients, which supports our results(Fang et al., 2020). Besides, the abundance of *Butyricicoccus* was found to be elevated in the intestines of infants with food allergies and was reported to be positively associated with allergy rhinitis(Ling et al., 2014, Zhu et al., 2020). In addition, we found that the relative abundance of the ranked first genus *Bacteroides* boosted significantly in Mild_AD patients. However, previous studies have reported a higher frequency of *Bacteroides* in eczema infants or a lower frequency of *Bacteroides* in eczema infants(Abrahamsson et al., 2012, Zheng et al., 2016). We speculate the inconsistent consequences may be caused by the different severity of AD patients.

Our results illustrated that the proteinogenic amino acid biosynthesis of the gut microbial community was limited, while pathways of purine nucleotide degradation metabolism were vigorous in AD patients. Interestingly, we found that the child pathways of vitamin B6 biosynthesis were severely deficient in people with severe AD. Previous studies have indicated that the microbiome in the intestine lumen converts pyridoxal phosphate into free vitamin B6, which enters the body fluid circulation through passive transport(Magnúsdóttir et al., 2015). Vitamin B6 deficiency is positively related to allergy. Lack of vitamin B6 can disrupt the body’s Th1-Th2 balance, leading to excessive Th2 reactions, contributing to allergies(Qian et al., 2017).

This study concentrated on the structural and functional alterations of adult AD patients’ gut microbiome. We have to point out that the clinical information such as allergic symptoms and gastrointestinal symptoms of the included samples were declared by the participants themselves, rather than strict medical diagnosis, which might cause the results to be inconsistent compared to previous studies. Besides, we performed amplicon sequencing on fecal samples, which was only accurate to the genus level and was difficult to reveal the gut flora composition at the species level and bacterial genome. Moreover, metabolites related to AD in our results were just predictions calculated by certain algorithms. Despite we found some evidence to support our results, verification of our predictions was required. Future studies with the strict medical diagnosis of allergies and gastrointestinal symptoms, and more clinical information of participants, such as dietary habits and medication status are warranted. Furthermore, whole genome shotgun sequencing and metabolomics sequencing of the gut microbiome should be carried out in the future, which can clarify the interaction between the gut microbiome and host in AD patients, to reveal potential targets and provide novel therapeutic strategies for AD.

In summary, the composition, structure, and functional metabolic pathways of the gut bacterial community of AD patients were distinct from those of healthy people but severe and mild AD patients had a similar overall structure of gut microbiome. We also discussed the possible interaction mechanism between the intestinal flora, functional metabolic pathways, and the host in AD patients. Gut microbiome dysbiosis and metabolic abnormalities may have essential implications for the pathogenesis of the gut-skin axis in AD patients. Further studies of metagenomics and metabolomics are warranted in order to investigate the relationship between the gut microbiome and atopic dermatitis and further develop beneficial methods of atopic dermatitis.

## MATERIALS & METHODS

### Study design and participants

Subjects were recruited from a community trial through the collaboration between The Chinese University of Hong Kong and the BioMed Technology Holdings Limited. A total of 259 participants’ fecal samples were collected and we excluded 25 of them who were less than the age of 18. Finally, a total of 234 people, distributed between the ages of 18-68, were included in this study, among which 130 participants with healthy skin were assigned to the normal group. After atopic dermatitis diagnosis and severity assessment using the scale of EASI by the dermatologist, there were 104 patients with atopic dermatitis (AD) including 53 mild AD patients and 51 severe AD patients. Additionally, 147 of 243 participants, including 78 AD patients and 60 healthy subjects provided the clinical information of allergy history, prevalence of diarrhea and constipation, BMI (body mass index) for subsequent patient characteristics analyses. Written informed consents were obtained from all subjects or their legal guardians.

### Sample collection, DNA extraction and 16S rRNA gene sequencing

Stool samples were homogenized in PurSafe® DNA and RNA preservative (Puritan, US) and subjected to beating with glass beads (425-600 μm, Sigma) for 1 hour by following the instructions provided. Microbial DNA was isolated from the stool samples with DNeasy Blood & Tissue Kit (Qiagen, Germany) according to the manufacturer’s instructions. The extracted DNA concentration of each sample was quantified using a Qubit™ dsDNA HS Assay Kit (Life technology) with Qubit 3 Fluorometer (Thermo Scientific). Amplicon library was constructed using 515F(5’-GTGCCAGCMGCCGCGG-3’)/907R(5’-CCGTCAATTTCMTTTRAGTTT- 3’) primer pair spanning targeting at V4-V5 hypervariable of 16S rRNA genes, together with adapter sequences, multiplex identifier tags, and library keys. 16S rRNA gene sequencing was performed using the Illumina MiSeq platform (Illumina, Inc., San Diego, CA) following the original Earth Microbiome Project Protocols. Finally, we obtained index barcodes and adapters removed pair-end clean reads for the downstream analysis(Caporaso et al., 2012).

### Microbiome data analysis

Microbiome bioinformatics analyses were performed using QIIME 2-2020.11, a plugin-based system, which integrates various microbiome analysis methods(Bolyen et al., 2019). Briefly, quality control and denoising filter of sequence data were processed by DADA2(Callahan et al., 2016) using the q2-dada2 plugin to obtain all observed amplicon sequence variants (ASVs)(Callahan et al., 2017). All ASVs were then aligned with mafft(Katoh et al., 2002) and used to generate a phylogenetic tree with fastree2(Price et al., 2010) for the downstream analyses via the q2-phylogeny plugin. Taxonomic assignment was performed using the q2-feature-classifier(Bokulich et al., 2018) plugin and a pre-trained Naive Bayes classifier which was based on SILVA v138 taxonomic reference database(Glöckner et al., 2017, Quast et al., 2012, Yilmaz et al., 2013).

### Functional profiling prediction with PICRUSt2

PICRUSt2 was developed to predict the metagenomic functional profiling of amplicon sequencing data(Douglas et al., 2020). We performed PIRCRUSt2 functions using the plugin of q2-picrust2 wrapped in QIIME 2-2019.7. LEfSe was carried out to identify the significantly enriched metagenome metabolic pathways across groups(Segata et al., 2011).

### Statistical analysi*s*

Statistical analysis was conducted in R 4.0.4. Diversity analyses were performed using the R package microeco (v0.3.2)(Liu et al., 2020). Alpha diversity was represented by the Shannon diversity index, Simpson diversity index, observed OTUs, Chao 1 richness index, abundance-based coverage estimator (ACE) index, and faith’s phylogenetic diversity index. Kruskal-Wallis rank sum test was performed to compare differences in alpha diversity across groups(Hill et al., 2003). Multivariate linear regression was carried out to adjust the clinical variables and batch effects. Beta diversity was calculated based on the Jaccard distance metric, Bray-Curtis distance metric, weighted UniFrac, and unweighted UniFrac distance metrics. The PERMANOVA test on beta diversity (999 permutations) was applied to compare the microbial community dissimilarity across groups using the adonis function in vegan R package to adjust the clinical variables and batch effects(Anderson, 2017). ANCOM, analysis of the composition of the microbiome, was performed to conduct differential abundance test across groups(Mandal et al., 2015). Linear discriminant analysis effect size (LEfSe) analysis was conducted to detect biomarker of the gut microbiome of each group with α equal to 0.05 and LDA score threshold of 2.0 using the online Galaxy platform (https://huttenhower.sph.harvard.edu/galaxy/)(Segata et al., 2011). Spearman correlation was carried out to estimate the correlation between the EASI score and the relative abundance of gut microbial signatures. Shapiro-Wilk normality test were carried out for normality of all data. Demographic characteristics across groups were compared using Wilcoxon rank-sum tests (two groups) or ANOVA (three groups) for continuous variables and Chi-square tests or the Fisher exact test for categorical variables. *P*-value < 0.05 represents statistical significance.

## Supporting information

Supplemental Tabels

## Data availability statement

The raw sequence data were deposited in the NCBI Sequence Read Archive with the accession number PRJNA778863.

## Contributor statement

SKWT and SKFL designed this study, obtained funding, recruited subjects, supervised microbiome analysis, provided clinical guidance, and prepare the manuscript. YW participated in the study design, performed bioinformatics analysis of microbiome data, interpreted the results, and wrote the manuscript. JH and LW assisted statistical analysis of microbiome data and interpreted the results. JCCT participated in the study design, recruited subjects, and collected fecal samples and clinical data. JZ, UKC, CJYL, and PLKS collected clinical samples, performed laboratory experiments, and conducted amplicon sequencing. All authors read and approved the final manuscript.

