## Supplemental Tabels for "Unique gut microbiome signatures among adult patients with moderate to severe atopic dermatitis in southern Chinese"

**Supplementary Tables**

**Table S1** Summary of alpha diversity analysis results of different grouping schemes

| Measure | 2 Groups | | 3 Groups | | | | | |
| --- | --- | --- | --- | --- | --- | --- | --- | --- |
|  | Normal VS AD | | Normal VS Mild AD | | Normal VS Severe AD | | Mild AD VS Severe AD | |
|  | *p* value | Sig. | *p* value | Sig. | *p* value | Sig. | *p* value | Sig. |
| Observed OTUs | 0.928 |  | 0.789 |  | 0.874 |  | 0.817 |  |
| Chao 1 | 0.859 |  | 0.701 |  | 0.878 |  | 0.777 |  |
| ACE | 0.766 |  | 0.551 |  | 0.834 |  | 0.673 |  |
| Shannon diversity index | 0.285 |  | 0.249 |  | 0.743 |  | 0.734 |  |
| InvSimpson index | 0.369 |  | 0.225 |  | 0.830 |  | 0.483 |  |
| Fisher | 0.901 |  | 0.799 |  | 0.933 |  | 0.876 |  |
| Faith pd | 0.844 |  | 0.831 |  | 0.792 |  | 0.842 |  |

Abbreviation: Sig., Significant

Sig. codes: * *p* < 0.05; ** *p* < 0.01; *** *p* < 0.001

**Table S2** Beta diversity analysis results based on four different metrics across groups

| Group | Metric | *p* value | Sig. |
| --- | --- | --- | --- |
| Normal VS AD | Jaccard distance metric | 0.001 | *** |
|  | Bray-Curtis distance metric | 0.001 | *** |
|  | Unweighted UniFrac distance metric | 0.025 | * |
|  | Weighted UniFrac distance metric | 0.181 |  |
| Normal VS Mild AD | Jaccard distance metric | 0.025 | * |
|  | Bray-Curtis distance metric | 0.030 | * |
|  | Unweighted UniFrac distance metric | 0.343 |  |
|  | Weighted UniFrac distance metric | 0.082 |  |
| Normal VS Severe AD | Jaccard distance metric | 0.001 | *** |
|  | Bray-Curtis distance metric | 0.001 | *** |
|  | Unweighted UniFrac distance metric | 0.031 | * |
|  | Weighted UniFrac distance metric | 0.429 |  |
| Mild AD VS Severe AD | Jaccard distance metric | 0.119 |  |
|  | Bray-Curtis distance metric | 0.103 |  |
|  | Unweighted UniFrac distance metric | 0.769 |  |
|  | Weighted UniFrac distance metric | 0.139 |  |

Abbreviation: Sig., Significant

Sig. codes: * *p* < 0.05; ** *p* < 0.01; *** *p* < 0.001

**Table S3** Relative abundance of species at the phylum level in AD and normal groups

| Kingdom | Phylum | AD (mean) | Normal (mean) | W value |
| --- | --- | --- | --- | --- |
| Bacteria | Firmicutes | 0.5738 | 0.5816 | 1 |
|  | Bacteroidota | 0.3243 | 0.3207 | 0 |
|  | Actinobacteriota | 0.0618 | 0.0546 | 0 |
|  | Proteobacteria | 0.0275 | 0.0244 | 0 |
|  | Fusobacteria | 0.0029 | 0.0074 | 0 |
|  | Desulfobacterota | 0.0046 | 0.0045 | 0 |
|  | Verrucomicrobiota | 0.0043 | 0.0041 | 0 |
|  | Cyanobacteria | 0.0002 | 0.0020 | 0 |
|  | Elusimicrobiota | 0 | 3.94e-06 | 0 |
|  | Synergistota | 0.0003 | 0.0003 | 0 |
|  | Unclassified | 5.67e-05 | 2.56e-05 | 0 |
|  | Campilobacterota | 2.96e-05 | 1.97e-05 | 0 |
| Archaea | Euryarchaeota | 0.0003 | 0.0004 | 0 |
|  | Halobacterota | 0 | 7.89e-06 | 0 |
|  | Thermoplasmatota | 3.45e-05 | 0 | 0 |

W value was calculated using ANCOM

**Table S4** Gut microbial biomarkers selected by LEfSe analysis

| Bacteria | Group | LDA score | *p* value |
| --- | --- | --- | --- |
| *g_Romboutsia* | Normal | 3.49 | 0.002 |
| *g_Clostridium_sensu_stricto_1* | Normal | 3.16 | 0.0008 |
| *f_Butyricicoccaceae* | Normal | 2.10 | 0.008 |
| *f_Erysipelotrichaceae* | Normal | 3.54 | 0.006 |
| *Blautia* | AD | 3.78 | 0.009 |
| *Butyricicoccus* | AD | 3.28 | 0.003 |
| *Lachnoclostridium* | AD | 3.22 | 0.045 |
| *Eubacterium_hallii_group* | AD | 3.12 | 0.015 |
| *Erysipelatoclostridium* | AD | 3.03 | 0.019 |
| *Megasphaera* | AD | 2.94 | 0.017 |
| *Oscillibacter* | AD | 2.87 | 0.019 |
| *Flavonifractor* | AD | 2.62 | 0.026 |
| *f_Oscillospiraceae* | AD | 2.93 | 0.001 |

**Table S5** Differentially enriched predicted functional pathways across groups using PICRUSt2 and LEfSe analysis.

| MetaCyc ID | MetaCyc pathway | Group | LDA score | *p* value |
| --- | --- | --- | --- | --- |
| PWY-4984 | Urea cycle | Normal | 2.00 | 0.001 |
| PWY490-3 | Nitrate reduction VI (assimilatory) | Normal | 2.08 | 0.045 |
| PWY-5659 | GDP-mannose biosynthesis | Normal | 2.13 | 0.01 |
| GLYCOLYSIS | Glycolysis I (from glucose 6-phosphate) | Normal | 2.14 | 0.038 |
| PWY-2941 | L-lysine biosynthesis II | Normal | 2.17 | 0.01 |
| PWY-6630 | Superpathway of L-tyrosine biosynthesis | Normal | 2.28 | 0.004 |
| PWY-6628 | Superpathway of L-phenylalanine biosynthesis | Normal | 2.29 | 0.004 |
| PWY-7315 | dTDP-N-acetylthomosamine biosynthesis | AD | 2.01 | 0.017 |
| PWY-7242 | D-fructuronate degradation | AD | 2.09 | 0.027 |
| P164-PWY | Purine nucleobases degradation I (anaerobic) | AD | 2.10 | 0.005 |
| PWY-1861 | Formaldehyde assimilation II (RuMP Cycle) | AD | 2.11 | 0.009 |
| GLYCOLYSIS-E-D | Superpathway of glycolysis and Entner-Doudoroff | AD | 2.12 | 0.035 |
| P441-PWY | Superpathway of N-acetylneuraminate degradation | AD | 2.12 | 0.044 |
| METH-ACETATE-PWY | Methanogenesis from acetate | AD | 2.16 | 0.001 |
| PWY-6608 | Guanosine nucleotides degradation III | AD | 2.24 | 0.004 |
| PWY-6353 | Purine nucleotides degradation II (aerobic) | AD | 2.32 | 0.0003 |
| SALVADEHYPOX-PWY | Adenosine nucleotides degradation II | AD | 2.37 | 0.0003 |
| PWY-5971 | Palmitate biosynthesis II (bacteria and plants) | AD | 2.38 | 0.027 |
| PRYIDOXSYN-PWY | Pyridoxal 5'-phosphate biosynthesis I | Mild_AD | 2.62 | 0.006 |
| PWY0-845 | Superpathway of pyridoxal 5'-phosphate biosynthesis and salvage | Mild_AD | 2.67 | 0.006 |
| PWY-6353 | Purine nucleotides degradation II (aerobic) | Severe_AD | 2.53 | 0.0002 |
| SALVADEHYPOX-PWY | Adenosine nucleotides degradation II | Severe_AD | 2.57 | 0.0002 |
