## Supplemental Figure Legends for "Unique gut microbiome signatures among adult patients with moderate to severe atopic dermatitis in southern Chinese"

**Supplementary Figures**

**Figure legends:**

**Figure S1** Boxplot of B/F ration across groups, *p* value was calculated using Kruskal-Wallis test.

**Figure S2** Bar plot of LEfse analysis results in the gut microbiome of Mild_AD, Severe_AD, and normal groups.
