## Supplementary figures and images for "Unique gut microbiome signatures among adult patients with moderate to severe atopic dermatitis in southern Chinese"

### Supplemental Figure 1

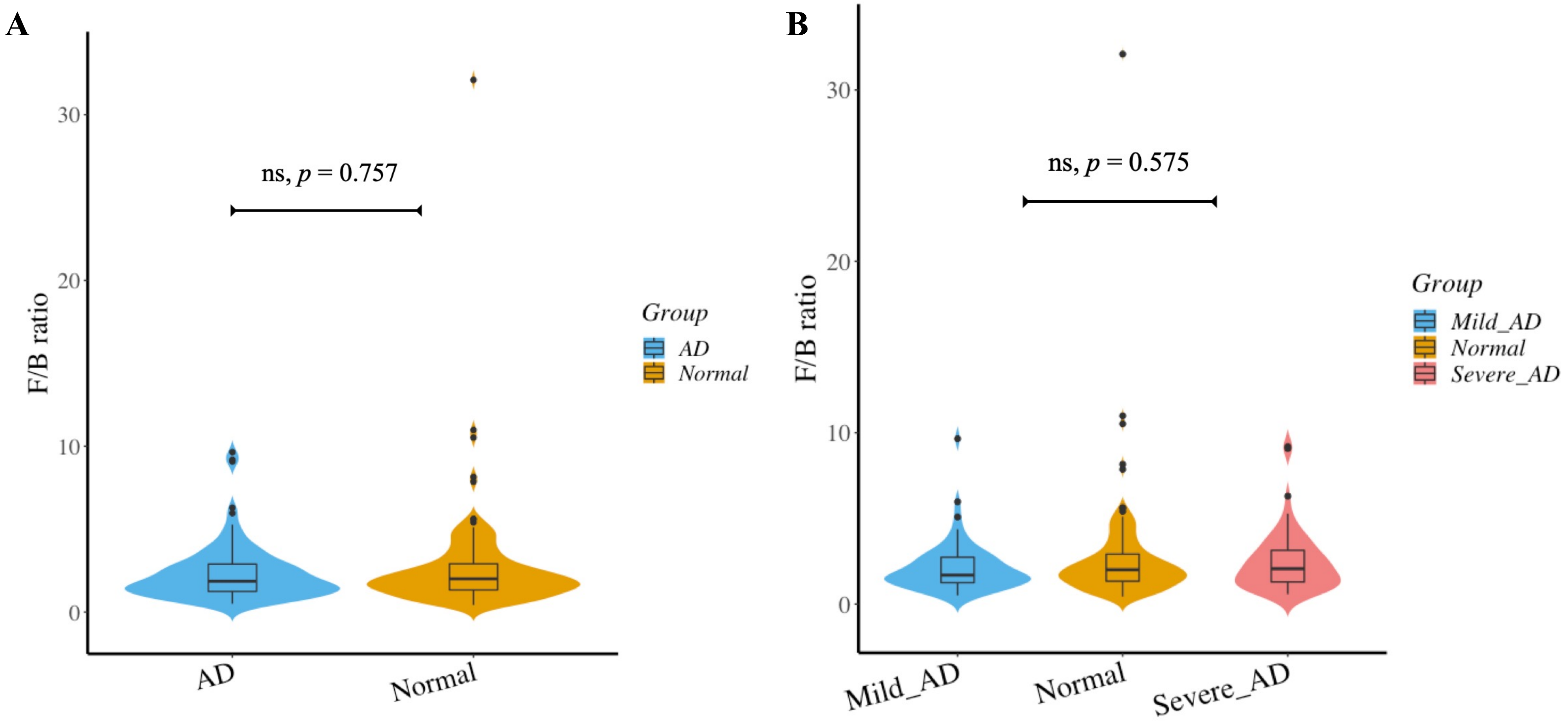

### Supplemental Figure 2

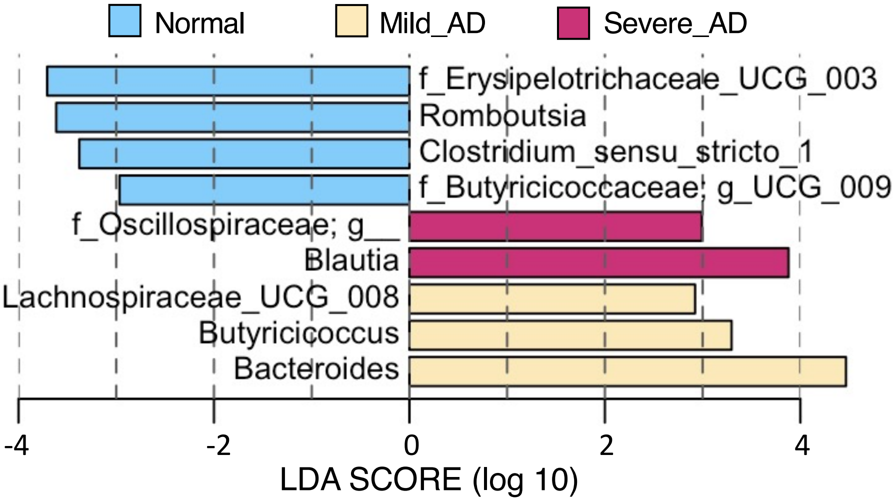
